# Sexual selection on body size, genitals and heterozygosity:effects of demography and habitat complexity

**DOI:** 10.1101/045724

**Authors:** Megan L. Head, Andrew T. Kahn, J. Scott Keogh, Michael D. Jennions

## Abstract

Environmental variation can maintain genetic variation in sexually selected traits if it affects the strength of directional selection. Specifically, environmental variation in sex-specific mortality will change the operational sex ratio (OSR), which predicts the intensity of mating competition. How the OSR affects selection for specific male traits is poorly understood; and it is unknown how often sexual selection is affected by interactions between the OSR and environmental factors that alter social variables such as mate encounter rates. Here, we experimentally manipulated the OSR and habitat complexity and quantified sexual selection on male mosquitofish (*Gambusia holbrooki*). In *G. holbrooki* there is high within-population variation in male size, which may exist because of a tradeoff between the ability to sneak copulate (favouring small males) and monopolize females (favouring large males). The success of each tactic is predicted to depend on the OSR, encounter rates and the ability to stealthily approach conspecifics. We show that, despite greater sharing of paternity under a male-biased OSR, neither the opportunity for selection, nor selection on male traits was affected by the OSR or habitat complexity. Instead, sexual selection consistently favored smaller males with high genetic heterozygosity (using >3000 SNP markers), and a relatively long gonopodium (intromittent organ).

## Introduction

Variation in the strength and form of sexual selection has generated much of the diversity in morphology, behavior and physiology between the sexes, across populations and among species (Andersson 1994; Pfennig and Pfennig 2010). Field studies have also shown that sexual selection can vary across populations and through time, be this between years (e.g., Chaine and Lyon 2008; Punzalan et al. 2010) or over a breeding season (e.g., Kasumovic et al. 2008; Milner et al. 2010; Wacker et al. 2014). Spatial and temporal variation in sexual selection is sometimes invoked to drive the evolution of discrete male mating types within a species (e.g., Gamble et al. 2003; Gray and McKinnon 2007; review: Buzatto et al. 2014). More generally, spatial variation in selection is thought to slow the erosion of genetic variation in sexual traits (Holman and Kokko 2014) that are usually under strong unidirectional selection (Hoekstra et al. 2001). Similar arguments have been made for the role of temporal variation in maintaining genetic diversity (Bell et al. 2010), although the evidence for temporal variation is mixed (Morrissey and Hadfield 2012; Siepielski et al. 2009). Explaining persistent phenotypic variation despite strong selection is a general challenge in evolutionary biology. Variation in sexual selection across heterogeneous environments is often posited as an explanation, but few studies directly address this question (Cornwallis and Uller 2010).

Despite evidence that sexual selection on focal traits varies within-species (i.e. among populations or between breeding cycles) we rarely understand why this occurs (Janicke et al. 2015). Most research on variation in sexual selection on focal traits comes from long-term observational studies of wild populations that draw inferences from annual or seasonal changes in plausible ecological factors (e.g., Kasumovic et al. 2008; Wacker et al. 2014). However, many ecological and demographic parameters tend to covary so it is difficult to identify specific parameters that drive variation in sexual selection. Similar problems arise when using population comparisons to identify environmental factors that affect sexual selection. For example, differences in male guppy (*Poecilia reticulata*) colouration are attributed to habitat differences in predation (e.g., Endler 1980), but predation also affects demography (e.g., Arendt et al. 2014; McKellar et al. 2009), which also can alter selection on male sexual signals. As always, experiments are required to determine which environmental factors have a causal effect. These are uncommon for field studies of naturally selected traits (see Morrissey and Hadfield 2012), and rare for field studies of sexually selected traits. Laboratory studies that manipulate environmental factors are commonplace but they are usually based on indirect measures of fitness (e.g., female association in two choice trials) rather than actual male reproductive success, and involve unnatural situations (e.g., small aquaria or tanks) (e.g., Seehausen and van Alphen 1998). Realistic experiments that measure actual male reproductive success are required.

Studies of variation in sexual selection often focus on the social environment, especially the operational sex ratio (OSR: the ratio of sexually receptive males to females) (review: Kvarnemo and Ahnesjo 1996). The role of the OSR in shaping sexual selection is debated (Klug et al. 2010; Krakauer et al. 2011). Variance in the reproductive success of the more common sex is expected to increase when the OSR is more biased due to greater competition for mates (Emlen and Oring 1977; Kvarnemo and Ahnesjo 1996; Shuster and Wade 2003). This is, however, not an inevitable outcome and depends on idiosyncratic features of each species (e.g., is harem defence easier or harder as the OSR becomes more biased?) (Jennions et al. 2012; Klug et al. 2010; meta-analysis: Moura and Cardoso Peixoto 2013). It is equally important to recall that not all variance in reproductive success is due to selection. Indeed, a positive relationship between the OSR and variance in reproductive success can arise simply due to a greater effect of stochasticity when the OSR is male-biased (Jennions et al. 2012). As importantly, the OSR might affect the various components of sexual selection in different ways so the observed effects depend on the type of mating system and which traits are measured (Kokko and Rankin 2006). For example, in a meta-analysis by Weir et al (Weir et al. 2011) it was noted that as the OSR becomes more biased the competitive sex becomes more aggressive (at least initially), but courts less. This is likely to lead to different estimates of how the OSR affects selection on courtship and fighting traits. Similarly, the effects of the OSR on mate choice, sperm competition and sexual coercion further complicate the relationship between the OSR and sexual selection (Fitze and Le Galliard 2008; Head et al. 2008). Finally, while rarely discussed, inconsistent effects of the OSR on the strength of sexual selection might arise because its effects are context-dependent and vary with other environmental factors (e.g., predation risk).

One ecological parameter that is of special interest is habitat complexity. Habitats vary in complexity over many spatial and temporal scales with profound effects on sexual selection (Myhre et al. 2013). For example, the transmission of mating signals depends on habitat complexity with open habitats generally allowing transmission of signals over greater distances. Such differences can affect how females perceive and assess male sexual signals and alter selection on males (e.g. sensory bias/drive (Boughman 2002; Endler and Basolo 1998). Habitat complexity can also affect selection on traits that affect male fighting success. For example, recent studies report higher aggression between male sticklebacks (*Gasterosteus aculeatus*) in open than complex habitats that might generate habitat-specific selection on body size (Lackey and Boughman 2013). Finally, the OSR and habitat complexity are both expected to alter mate encounter rates (Myhre et al. 2013), and habitat complexity can generate variation in local OSRs and densities (i.e., those directly experienced by each female). Such effects could alter the levels of both male-male competition and female choice. For example, in guppies greater habitat complexity reduced interference competition between males and increased female mating receptivity (Hibler and Houde 2006).

Here we conduct a manipulative experiment to investigate how habitat complexity and the adult sex ratio (ASR), hence OSR, interact to affect sexual selection on male mosquitofish (*Gambusia holbrooki).* This species is well suited to test how these factors affect sexual selection because it inhabits a range of habitats varying in structural complexity (from streams and ponds to lakes) and sex ratios vary predictably over the breeding season (Kahn et al. 2013). Furthermore, past studies have identified target traits that might be under sexual selection (Head et al. 2015; McPeek 1992; Pilastro et al. 1997). The mating system is characterised by sexual coercion, whereby males incessantly attempt to mate by approaching females from behind and thrust their gonopodium into her gonopore (Bisazza 1993; Bisazza and Marin 1995). This generates several, sometimes conflicting, modes of sexual selection on males. First, male-male competition favours large males that dominate access to females (Bisazza and Marin 1991). Second, female mate choice favors larger males with a longer gonopodium (Bisazza et al. 2001; Head et al. 2015; Kahn et al. 2010; McPeek 1992). Third, sexual coercion favors smaller males who are more adept at sneaking copulations (Pilastro et al. 2003; Pilastro et al. 1997). Four, post-copulatory mechanisms are likely to affect male fertilization success under sperm competition (Evans et al. 2003a). Five, it has recently been shown that heterozygosity is positively correlated with male reproductive success (XXXX, in prep). All of these selective mechanisms contribute to male reproductive success and could be influenced by both the OSR and habitat complexity. Finally, because these fish are small (15-45mm) and naturally breed in small water bodies it is possible to create semi-natural breeding conditions using artificial ponds.

Many studies that investigate the effects of the OSR on sexual selection focus on how it influences variance in reproductive success and the *opportunity* for selection (*I*), but *I* is not necessarily a good index of the actual *strength* of selection on phenotypic traits (Jennions et al. 2012; Kokko et al. 2012). Its reliability is an empirical question that is likely to vary among species and depend on the specific male traits and environments being examined. We therefore compared the value of *I* in different environments with how selection acted on three focal male traits: male body size, relative gonopodium length and genome wide heterozygosity.

## Methods

### Experimental design

We independently manipulated the adult sex ratio (ASR), which is the same as OSR if the OSR is calculated at the start of the experiment, and habitat complexity in ponds (1m diameter, 15cm depth) in a greenhouse using a 2 × 2 factorial design. We had two levels of habitat complexity. In the ‘simple’ habitat the pond floor was lined with gravel, the pond walls were lined with white plastic, and there was no vegetation or cover. The ‘complex’ habitat was the same, but we added a crossed network of white plastic baffles to create a series of interconnected compartments (Fig. S1). Manipulating habitat complexity in this way ensured that the manipulation was applied evenly across the whole pond. This manipulation of habitat complexity is similar to that used in a previous study (Hibler & Houde 2006) which found that increased visual isolation in complex habitats altered sexual behaviour in guppies. We also had two levels of ASR. The female-biased ASR consisted of 10 males and 20 females in a pond. The male-biased ASR consisted of 10 males and 5 females in half a pond. We avoid confounding changes in fish density and the number of males by adjusting the pond size and keeping the number of males constant. Fish density across all treatments was 0.25 fish per litre, which falls within natural fish densities seen in the wild (e.g. Jordan et al 1998) and those used in previous studies of sexual behaviour in poeciliid fishes (e.g. Marriette et al 2010; Magurran & Seghers 1995; Devigili et al 2015; Hibler & Houde 2006). Any treatment differences are therefore due to the ASR, and not covarying demographic parameters (see Head et al. 2008). We set up six blocks (one replicate/ treatment) (*n*=6x4x10=240 males).

### Experimental protocol

We used *G. holbrooki* from ponds in Canberra, Australia (35°14’27”S, 149°5’27”E and 35°14”13”S, 149 ° 5’55”E). These ponds are less than 2km apart and likely to be connected during periods of high rainfall. Experimental males were caught from the wild. Experimental females were lab reared offspring of wild caught females. This ensured females were virgins at the beginning of our experiment. We collected females from the wild and allowed them to give birth. The fry were then placed in 3l aquaria in groups of up to five. From 4 weeks of age onward these fry were checked weekly for signs of maturation. As soon as we could determine their sex (elongation of the anal fin for males, development of eggs visible through the body wall for females) fish were placed in single sex tanks. Virgin females were 3-9 months old before being used in our experiment. The use of virgin females ensured that all offspring were sired by males from our experimental ponds. We could not use wild caught females as they store sperm (Pyke 2005).

Importantly, prior to placement in experimental ponds, both sexes underwent a priming period. This mimicked the experimental conditions that fish would later experience to ensure that paternity results reflected the treatments experienced and not the change from stock to experimental conditions. For priming, focal males and females were placed in experimental ponds with the appropriate number of individuals of the opposite sex. Focal males were placed with stock females, and focal females were placed with stock males whose gonopodium tip had been removed to prevent sperm transfer (Mautz 2011). Following 4 days of priming, focal fish were placed directly into their respective experimental treatments, and stock fish were returned to stock tanks.

Once in experimental ponds focal fish had 14 days to interact and mate. They were fed thawed frozen *Artemia* nauplii twice daily. The female-biased treatments were fed twice the amount of food as the male-biased treatments since there were twice as many fish.

Males were euthenised after being removed from the experimental ponds. We photographed their left side alongside a scale using a digital camera (Nikon Coolpix 5700) mounted to a dissecting microscope (Leica Wild MZ8). We later measured male standard length and gonopodium length in *Image J.* Males were then preserved in absolute ethanol and stored at ‐20°C.

Once females were removed from the experimental ponds they were anaesthetised in ice slurry, photographed (see above) and then placed individually in 1l tanks. Each tank contained a gravel substrate, plastic aquarium plants and a mesh divider to reduce maternal cannibalism. Tanks were checked twice daily for fry until the female had either produced two broods or three months had passed. Females were kept at 27°C±1°C on a 14:10 light:dark cycle and fed live *Artemia* twice daily. When a female gave birth she was placed in a new 1l tank if it was her first brood. If it was her second brood she was euthenised and preserved for genotyping. Fry were euthanized (< 24h after birth), and preserved in family groups of up to 10 fry/vial.

### Sampling for paternity analysis

To determine male reproductive success we took tissue samples from up to five mothers (on average 4.1 per pond, *n* = 100 in total), all possible sires (10 per pond, *n* = 240 in total) and all offspring (mean: 35.2 per pond, *n* = 844 in total) from the selected mothers for each pond. In the male-biased ASR treatment we therefore sampled all mothers that gave birth and in the female-biased ASR treatment we randomly sampled five females that gave birth. DNA was extracted from the tail muscle/caudal fin for adults and from the whole body (excluding head) for fry, using Qiagen DNeasy Blood and Tissue Kits (Qiagen, Victoria, Australia) following the manufacturer’s instructions.

After extraction, DNA samples were sent to a commercial genotyping service - Diversity Arrays. The details of the process are described in the Supplementary Materials. We obtained a data set of approximately 3171 SNPs with an average call rate of 97.7% and a reproducibility rate of 99.3%. From the selected SNPs we calculated a Hamming Distance Matrix of all 1185 individuals (potential sires, mothers, and offspring) to determine paternity. Recent studies show that as few as 30 optimized SNPs are sufficient to differentiate among 100,000 individuals using Hamming Distance values (HDV) (Hu et al. 2015). All fry were lined up against their mother and siblings and the HDVs evaluated to cross check for any sample mix ups. None were detected. HDVs were then compared against each of the 10 potential sires. The sire/fry with the lowest value was considered a match. We could assign paternities unambiguously for all 844 fry.

### Heterozygosity

Our calculation of heterozygosity is simply the number of SNP markers that were scored as heterozygous divided by the total number of successfully classified markers for that fish.

### Data analysis

#### Opportunity for Selection

We calculated the opportunity for selection (*I*) for males within each pond as the variance in the total number of offspring sired divided by the mean number of offspring sired per male (Shuster and Wade 2003). This calculation was based on paternity data from the five genotyped broods per pond. We ran a general linear model to test the effects of habitat complexity, ASR and their interaction on *I*_*s*_. The response variable was transformed using the power transform function in the “car” package of R. This model gave the same results as a generalised linear model with untransformed data and quasipoisson error structure, but the latter had a poorer fit. We also tested whether the ASR and habitat complexity influenced how many males shared paternity in each brood. We ran a generalised linear mixed model (GLMM) with the number of sires per brood as the response variable and ASR, habitat and their interaction as fixed effects. Pond identity was treated as a random effect. This model gave qualitatively similar results to a linear mixed model on transformed data.

#### Sexual selection on males

To determine which male traits influenced his reproductive success and whether this varied across different socio-environmental context we ran a GLMM. We treated the number of offspring each male sired as the response variable. ASR and habitat complexity were specified as fixed factors. Male standard length (logged), relative gonopodium length (residuals of the regression of log gonopodium length on log male length) and heterozygosity were included as covariates. Interactions between each of the three male traits and the two experimental factors were included in the model. We treated pond as a random effect and specified a Poisson error structure. To account for overdispersion we included individual as a random effect (Harrison 2014). Following this correction our data was underdispersed (dispersion parameter = 0.0131) and thus conservative. An analysis using a power transformation of the dependent variable and Gaussian error structure gave qualitatively similar results.

Neither the ASR or habitat complexity influenced the relationship between the number of offspring sired and any of the male traits (see *Results*) so we calculated experiment wide selection gradients using a linear multiple regression (Lande and Arnold 1983). We treated the relative number of offspring a male sired (calculated within ponds) as the response variable and log male length, relative gonopodium length and heterozygosity as predictor variables. All predictor variables were standardised across the experiment (mean = 0, s.d. = 1). Standardising traits within ponds gave very similar selection gradient estimates. Significance values were obtained from the same model except that the relative number of offspring sired was power transformed to account for its non-normal distribution and pond identity was treated as a random effect.

#### Reproductive success of females

To determine whether our treatments influenced female reproductive output we ran GLMMs. The number of broods (0, 1 or 2) was analysed using an ordinal logistic regression in the package “ordinal” using the command clmm to allow for random effects. For those females that did have offspring we also analysed the number of offspring in a female’s first brood, gestation time (days between leaving the pond and giving birth), and the total number of offspring a female produced using models with Poisson error that included an individual level random effect when data were over-dispersed (Harrison 2014). In all models ASR, habitat and female standard length (centred to a mean of 0; (Gelman 2008)) and their interactions were specified as fixed effects. Pond identity was treated as a random effect.

In all analyses, including block as a random effect did not influence our results. For simplicity this term were excluded from our models (Bolker et al. (2009) does not recommend including random factors with fewer than 5-6 levels). All analyses were conducted in R version 3.2.0 (Team 2015).

## Results

### The opportunity for selection on males

Neither the adult sex ratio (estimate ± SE = ‐1.682 ± 1.140, t = ‐1.476, p = 0.156), habitat complexity (estimate ± SE = ‐0.650 ± 1.140, t = ‐0.570, p = 0.575) nor the interaction between them (estimate ± SE = 0.600 ± 1.612, t = 0.372, p = 0.714) influenced the opportunity for selection on males. The mean number of sires per brood was greater under a male-biased than female-biased ASR (estimate ± SE = 0.420 ±0.192, Z = 2.181, p = 0.029), but it did not depend on habitat complexity (estimate ± SE = ‐0.095 ±0.195, Z = 0.488, p = 0.626), nor was there an interaction between ASR and habitat complexity (estimate ± SE = ‐0.125 ± 0.280 Z = 0.446, p = 0.655) (Fig. 1). This finding was not confounded by female fecundity depending on the ASR (see below).

**Figure 1.**
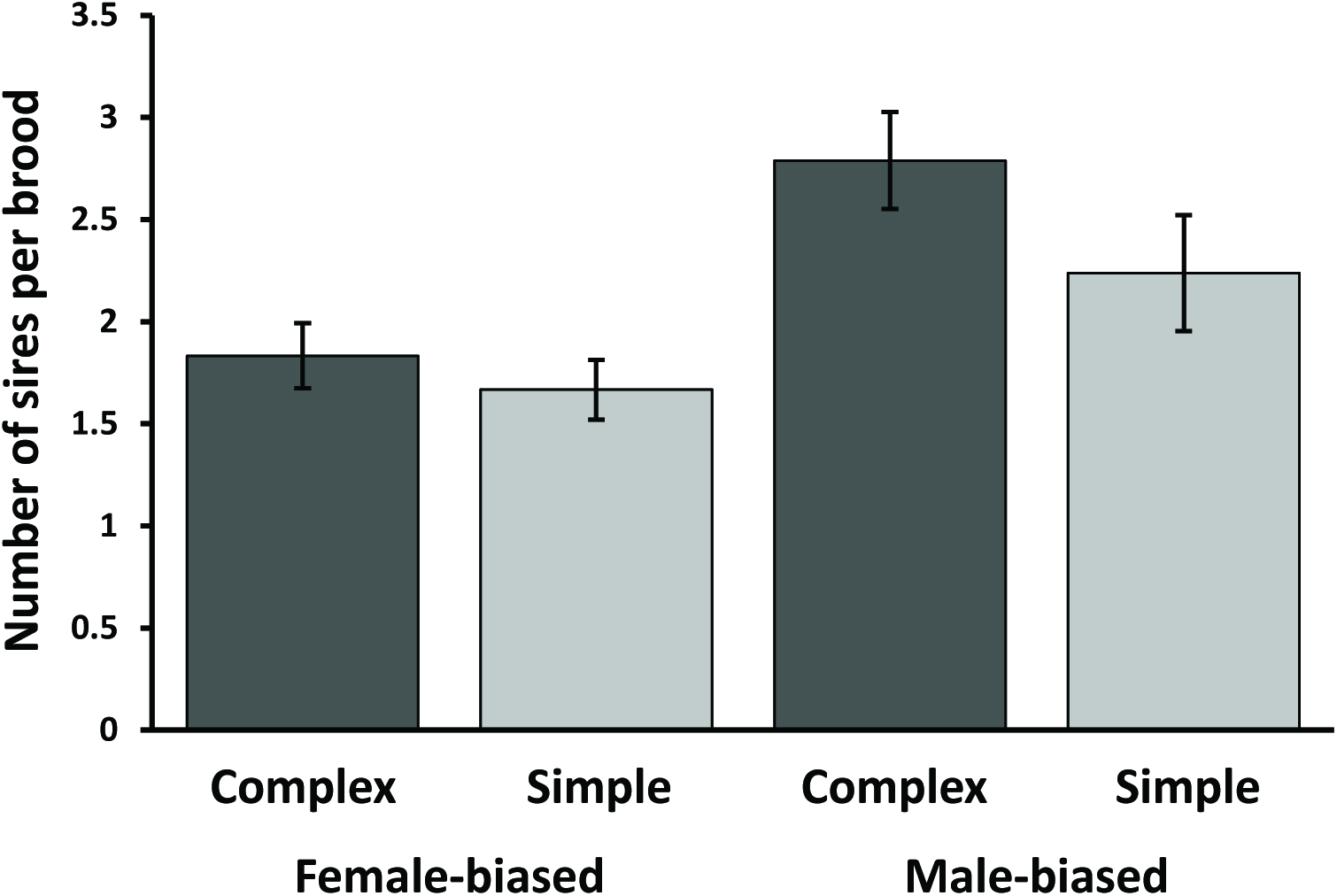
The mean number of sires contributing to each brood (±SE) within each treatment.

### Sexual selection on males

More heterozygous males were smaller (r=-0.164, t_(234)_= 1.354, p=0.012), and had a relatively longer gonopodium for their body size (r=0.187, t_(232)_= 2.901, p=0.004). Smaller males and more heterozygous males both had significantly greater reproductive success, but there was no effect of the ASR or habitat complexity on sexual selection on male traits (Table 1). When we examined net selection across all four treatments, the selection gradients were significant for all three male traits, as they were each independently correlated with reproductive success (Table 2). In 22 out of 24 ponds election favored more heterozygous males, in 19 out of 24 ponds selection favored smaller males, and in 20 out of 24 ponds selection favored males with a relatively long gonopodium.

**Table 1.** The effects of ASR and habitat complexity on the relationship between male traits and the number of offspring sired.

| Trait | Term | Estimate | SE | z | P |
| --- | --- | --- | --- | --- | --- |
| Number of Offspring | Intercept | 16.2477 | 11.859 | 1.370 | 0.171 |
|  | ASR (m) | -2.989 | 15.638 | -0.191 | 0.848 |
|  | Habitat (s) | 0.259 | 16.269 | 0.016 | 0.987 |
|  | Log Standard length (SL) | -18.502 | 7.924 | -2.335 | **0.020** |
|  | Relative gonopodium length (Gono) | 0.471 | 0.332 | 1.420 | 0.156 |
|  | Percent heterozygosity (Het) | 31.359 | 10.208 | 3.072 | **0.002** |
|  | ASR(m)*Habitat (s) | 1.971 | 21.350 | 0.092 | 0.926 |
|  | ASR(m)*SL | 1.213 | 10.427 | 0.116 | 0.907 |
|  | ASR(m)*Gono | -0.358 | 0.508 | -0.704 | 0.482 |
|  | ASR(m)*Het | 4.544 | 14.732 | 0.308 | 0.758 |
|  | Habitat(s)*SL | 2.071 | 10.895 | 0.190 | 0.849 |
|  | Habitat(s)*Gono | 0.680 | 0.506 | 1.346 | 0.178 |
|  | Habitat(s)*Het | -11.301 | 13.941 | -0.811 | 0.418 |
|  | ASR(m)*Habitat(s)*SL | 3.009 | 14.311 | 0.210 | 0.833 |
|  | ASR(m)*Habitat(s)*Gono | 0.251 | 0.755 | 0.333 | 0.739 |
|  | ASR(m)*Habitat(s)*Het | -21.053 | 19.285 | -1.092 | 0.275 |

Bold indicates significant effects

**Table 2.** The vector of experiment wide standardised linear selection gradients (β) for male traits in *Gambusia holbrooki.* Relative fitness calculated was calculated within ponds and male traits were standardised across the experiment. Selection gradients were estimated using linear multiple regression and the significance of these gradients was determined using a linear mixed model with power transformed relative fitness as the response variable to account for non-normal distribution of the data and pond included as a random effect to account for potential non-independence of data from the same pond.

| Trait | B (SE) | p |
| --- | --- | --- |
| Percent heterozygosity | 0.355 (0.103) | **<0.001** |
| Log standard length | -0.205 (0.101) | **<0.001** |
| Relative gonopodium length | 0.348 (0.101) | **<0.001** |

Bold indicates significant effect

### Reproductive success of females

Neither gestation time, the number of offspring in the first brood nor the total number of offspring a female produced were influenced by the adult sex ratio, habitat complexity or the interaction between them (Table 3). The number of broods per female was, however, influenced by both a female’s length and the ASR. With a female-biased adult sex ratio smaller females produced more broods than larger females, whereas with a male-biased sex ratio there was a weak relationship in the opposite direction. There was no effect of habitat complexity. Trait means for each treatment are shown in Table 4.

**Table 3.**
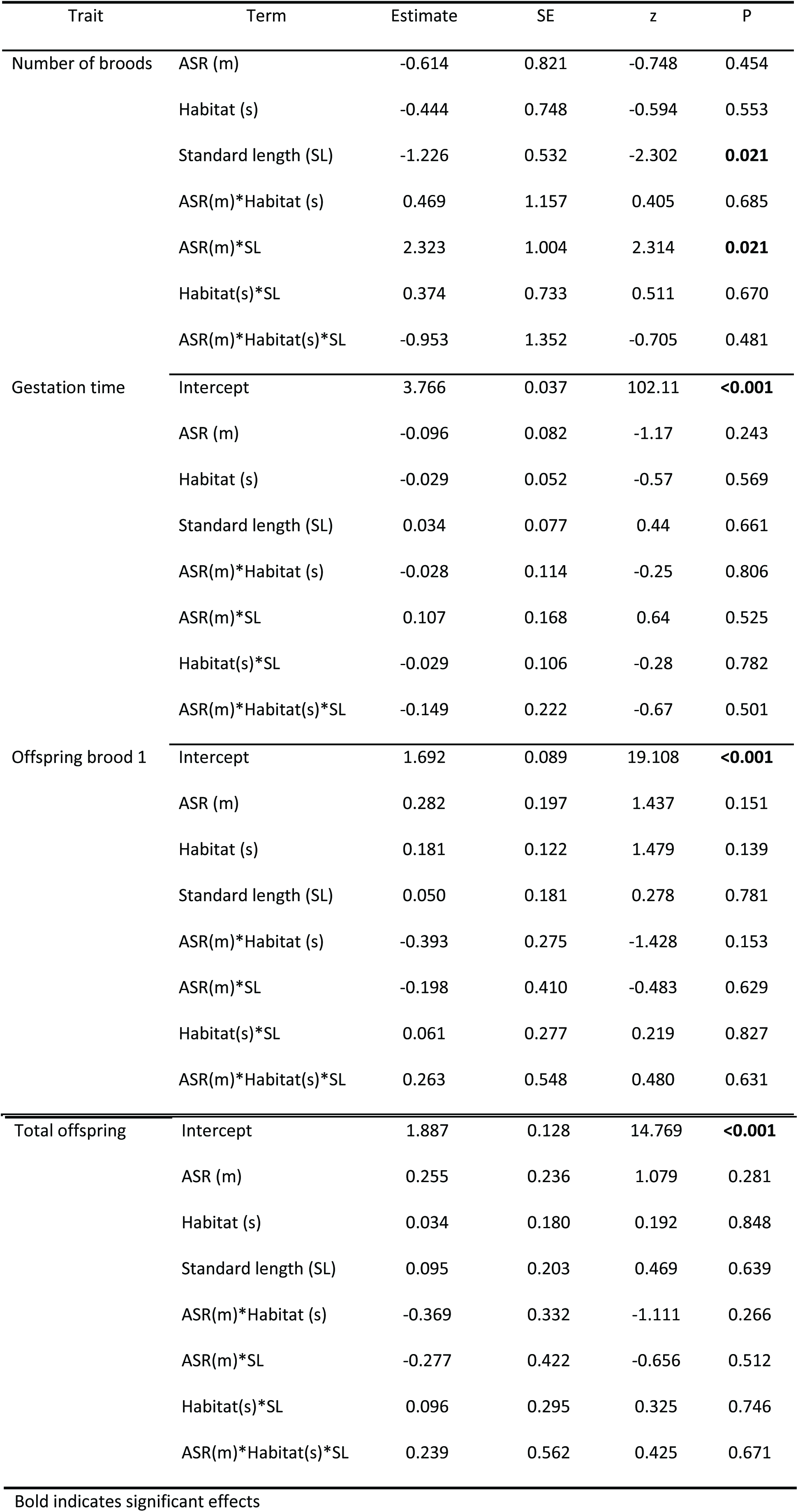
The effects of ASR and habitat complexity on female reproductive output.

**Table 4.**
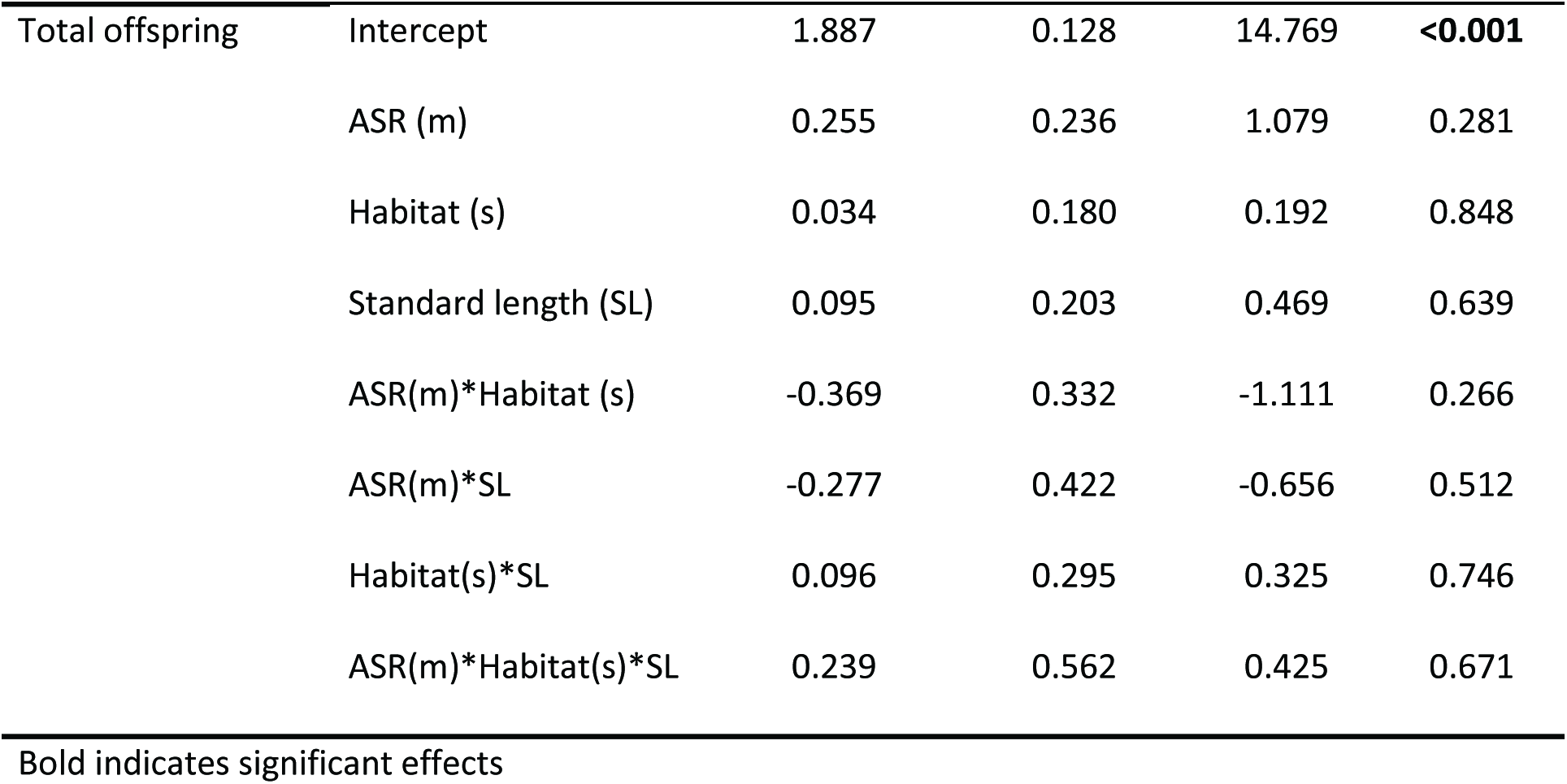
Female reproductive output in each treatment, mean (SE)

## DISCUSSION

Spatial and temporal environmental heterogeneity is often invoked as a factor helping to maintain variation in traits that are under directional selection (Cornwallis and Uller 2010; Gray and McKinnon 2007). We experimentally tested how two key ecological and demographic parameters - the adult sex ratio (ASR; which is equivalent to the OSR at the start of the experiment) and habitat complexity - influence sexual selection on male mosquitofish *Gambusia holbrooki.* The opportunity for selection (*I*) on males was constant across environments but, more importantly, our estimates of sexual selection on focal male traits did not differ across environments. Small males, males with a relatively larger gonopodium, and more heterozygous males had greater reproductive success in all cases. Our results suggest that altering mate encounter rates has little effect on sexual selection in *G. holbrooki.*

### The adult or operational sex ratio

Male-biased sex ratios lead to greater competition for mates, and it is usually assumed that this will also increase variation in male mating and reproductive success (Emlen and Oring 1977; Shuster and Wade 2003). Instead we found that the mean number of sires per brood became greater with a more male-biased sex ratio. All else being equal, a greater sharing of paternity should lower *I* because it reduces variation in male reproductive success. However, the adult sex ratio had no detectable effect on I. This suggests that even though more sires contributed to each brood this did not affect the distribution of paternity among males. This is expected if there is consistency among females in which type of males contribute to paternity. Our results agree with some studies that use variance based indices to measure the potential/opportunity for selection (e.g., Head et al. 2008; Mills et al. 2007) but contrast with others (Jones et al. 2004; Klemme et al. 2007) (for a review of indices see: Henshaw et al. in press).

There was no evidence that the sex ratio influenced selection on male traits in *G. holbrooki.* Numerous studies have shown that the ASR or OSR (in many experimental studies the two are interchangeable: see Kokko and Jennions (2008)) affect mating behaviour (e.g., Bretman et al. 2012; Holveck et al. 2015) and indices of variance in male reproductive success (Henshaw et al. in press). However, few studies experimentally manipulate the ASR to test whether it affects sexual selection on specific male traits. This is surprising as the evolution of male traits depends on how they relate to fitness (i.e., number of offspring sired). Of the experimental studies that have looked at how selection resulting from variation in reproductive success acts on male traits, the results are mixed. In guppies (*Poecilia reticulata,* Head et al. 2008) and bank voles (*Clethrionomys glareolus,* Mills et al. 2007) there was no effect of the ASR on the strength of selection on male sexual traits. In contrast, in two spotted gobies (*Gobiusculus flavescensa,* Wacker et al. 2013) and rough skinned newts (*Taricha granulosa,* Jones et al. 2004) selection on male traits was stronger when the ASR was more male-biased. Finally, and contrary to some expectations, selection on male traits was weaker when the ASR was more male-biased in studies of bank voles (*Clethrionomys glareolus,* Klemme et al. 2007) and common lizards (*Lacerta vivipara,* Fitze and Le Galliard 2011).

The conflicting results in previous studies of how the ASR affects sexual selection might be partly due to the confounding effects of other ecological parameters, especially those that determine how many individuals interact at any given moment (e.g., due to habitat complexity, or factors that influence population density (see: Kokko and Rankin 2006)). However, in *G. holbrooki* there was no evidence that habitat complexity affected sexual selection on the measured traits. It has generally been argued that whether the OSR affects sexual selection will depend on the mating system (e.g., Kokko and Rankin 2006). Of the aforementioned studies that tested whether selection on male traits is ASR-dependent, breeding in *G. holbrooki* is most similar to that in guppies, *Poecilia reticulata* where Head et al (2008) also found no effect of the ASR on sexual selection. In both these poeciliid fishes males increase their reproductive success through forced copulations (‘sneak mating’) and there is post-copulatory sexual selection due to sperm competition (Evans and Magurran 2001) and cryptic female choice (Evans et al. 2003b; Gasparini and Pilastro 2011). Post-copulatory sexual selection might dampen potential effects of the ASR on sexual selection if there is a trade-off between traits associated with elevated pre‐ and post-copulatory sexual selection (see (Devigili et al. 2015).

### Habitat complexity and sexual selection

Habitat complexity did not mediate the effect of the ASR on sexual selection, nor did it directly influence selection on male traits, even though habitat complexity alters sexual behaviour in other poeciliid fishes (e.g., Hibler and Houde 2006). Furthermore, habitat variation is important in shaping sexual traits in many species, which is why ecological factors are often implicated in maintaining population variation in sexual traits (Cornwallis and Uller 2010) and even in speciation (Maan and Seehausen 2011). For example, habitat differences in gravel size can lead to divergence in male colouration in guppies (*Poecilia reticulata,* Endler 1980). Similarly, variation in habitat complexity affects selection on male advertisement call in cricket frogs (*Acris crepitans,* Ryan et al. 1990). The difference between these studies and ours might reflect the types of traits being measured.

Colouration and vocalisations are sexual signals whose transmission and detection is habitat-dependent. In contrast, the traits we measured such as body size, relative gonopodium size and heterozgosity are integrally related to male quality. Here a more likely mechanism by which habitat complexity would alter selection is via effects on demography and mate encounter rates, which then affect how females assess males or shift the balance between different modes of sexual selection (e.g. mate choice vs coercion).

### Traits under sexual selection in *G. holbrooki*

We detected strong directional selection on males for body size, relative gonopodium length and heterozygosity. We consider each trait in turn.

Smaller male *G. holbrooki* had greater reproductive success. This has long been assumed to be the case in *Gambusia spp* based on behavioural evidence for insemination success (e.g., Pilastro et al. 1997), but actual paternity data has been lacking (although Deaton (2008) found a large male advantage based on paternity analysis of 27 trials where a small and a large male competed freely for access to a female within small aquaria). As in many species of poeciliid fishes, male mosquitofish vary substantially in size (range in this experiment 19 – 32mm). Understanding how this variation persists despite strong directional selection is a major challenge in evolutionary biology (Barton and Turelli 1989). Potential explanations include context-dependent selection (Cornwallis and Uller 2010), genic capture (Tomkins et al. 2004), fitness trade-offs between traits (Blows et al. 2003), and trade-offs between the effect of a given trait under different modes of selection (Devigili et al. 2015; Johnston et al. 2013). We could not identify an ecological factor maintaining variation in male size in *G. holbrooki,* but our results suggest that it is not due to variation in habitat complexity, nor to ecological factors that affect the ASR.

Male *G. holbrooki* with a relatively long gonopodium for their body size had higher reproductive success. Similar positive directional selection on gonopodium length has been shown previously in guppies, *P. reticulata* (Devigili et al. 2015; Evans et al. 2011). This could be due to female choice for males with a long gonopodium (Kahn et al. 2010; Langerhans et al. 2005), or a greater ability to inseminate females coercively (Evans et al. 2011). Interestingly, XXXX et al (in review) recently showed no increase in reproductive success for males from lines artificially selected for greater relative gonopodium length. This suggests that although relative gonopodium length is heritable (i.e., it evolved under artificial selection) and there is directional selection for males with a relatively long gonopodium, this might not be due to selection of a relatively long gonopodium (see Morrissey (2014) for a discussion on the distinction between ‘selection for’ and ‘selection of’ a trait). That is, an unmeasured factor might cause both greater relative gonopodium length and higher reproductive success. A likely candidate is body condition (see also Kruuk et al. 2002). This is a reminder of the easily overlooked fact that estimates of selection gradients can only truly estimate direct selection on traits if all relevant covarying traits are measured (Lande and Arnold 1983).

Male *G. holbrooki* with a higher heterozygosity had greater reproductive success. This is a finding that we have since replicated in a second paternity analysis study using an experimental design which systematically manipulates heterozygosity (XXXX in prep). Studies of heterozygosity fitness correlations (HFCs) show that homozygosity negatively affects fitness-enhancing traits (reviews: Chapman et al. 2009; Coltman and Slate 2003; Szulkin et al. 2010). There are, however, relatively few HFC studies that link heterozygosity to male reproductive success under sexual selection (i.e. control for male mortality). Of these, several studies show that lower heterozygosity decreases male reproductive success (e.g., water dragons, *Intellagama lesueurii* (Frere et al. 2015); Black rhinoceros, *Diceros bicornis michaeli* (Cain et al. 2014); zebra finches, *Taeniopygia guttata* (Forstmeier et al. 2012); house mice, *Mus musculus musculus* (Thoss et al. 2011); blue tits, *Cyanistes caeruleus* (Olano-Marin et al. 2011)), although this is not always true (e.g., Great tits, *Parus major* (Chapman and Sheldon 2011)). We observed a strong positive relationship between heterozygosity and male reproductive success (*r*=0.267) compared to a mean value for HFCs of *r*=0.05 (meta-analysis: Chapman et al 2009). There are several reasons why we see a strong relationship in *G. holbrooki.*

First, we had a better estimate of genome wide heterozygosity (Balloux et al. 2004). Microsatellite markers are generally 4-10 times more variable than SNPs (Mariette et al. 2002; Morin et al. 2004). Even so, the 3171 SNP markers we used is equivalent to using over 300 microsatellite markers. To date, most HFC studies use fewer than 20 microsatellite markers (Chapman et al. 2009). Second, traits that are more closely related to actual fitness are more likely to suffer inbreeding depression (Kristensen et al. 2010). The studies in the meta-analysis of Chapman et al. (2009) mainly report HFC correlations for morphological, physiological and life-history traits; very few studies provide direct fitness estimates such as reproductive success. As such, the average HFC in Chapman et al. (2009) is likely to be an underestimate of the true link with fitness (Chapman and Sheldon 2011). Third, the HFC is likely to depend on a population’s demographic history, with relationships being weaker in highly outbred or inbred populations where variation in heterozygosity is lower. Relatively low levels of heterozygosity (although still within the range of natural populations (Vera et al 2016), and substantial variation in heterozygosity (17-36%) that we see in our study may be particularly conducive to detecting inbreeding depression. Interestingly, another recent study that used a large number of SNPs to estimate inbreeding also found strong inbreeding depression when looking at fitness traits in red deer (*Cervus elaphus*) (including lifetime reproductive success) even though variation in heterozygosity was relatively low (Huisman et al in press).

There are many mechanisms whereby lower heterozygosity could reduce fitness. Previous studies have shown that inbred (i.e., more homozygous) males can be less attractive as mates (review:

Pusey and Wolf 1996), and sometimes produce less competitive ejaculates (e.g., Michalczyk et al. 2010; Simmons 2011; but see XXXX in review). Alternatively, low heterozygosity/inbreeding could affect traits such as locomotion (Kjaersgaard et al. 2014; Manenti et al. 2015) or cognition (Fareed and Afzal 2014) that reduce a male’s ability to locate and/or inseminate females. Regardless of the mechanism, however, a strong effect of heterozygosity on paternity is likely to have wider implications. For example, it is could select for female avoidance of inbred males which could affect the persistence and recovery of small populations (Keller and Waller 2002), because it might alter the ease with which genetic variation persists (i.e., fewer sires reduces genetic diversity, but mating with more genetically diverse males increases genetic diversity).

#### Conclusion

We found that sexual selection in the mosquitofish was consistent across habitat types that differed in two ecological parameters that affect mate encounter rates. While persistent ecological differences between habitats is clearly important for generating divergence between many species (reviewed in: Maan and Seehausen 2011), the role of temporal and spatial habitat variation in generating variation within species is less well understood (Cornwallis and Uller 2010). Experimental studies like the one described here looking at sexual selection in different environments and over different time/spatial scales are needed if we are to understand how ecological variation contributes to altering the strength and form of sexual selection and how organisms are likely to respond to changing environments.

## Supplementary methods

**Fig. S1.**
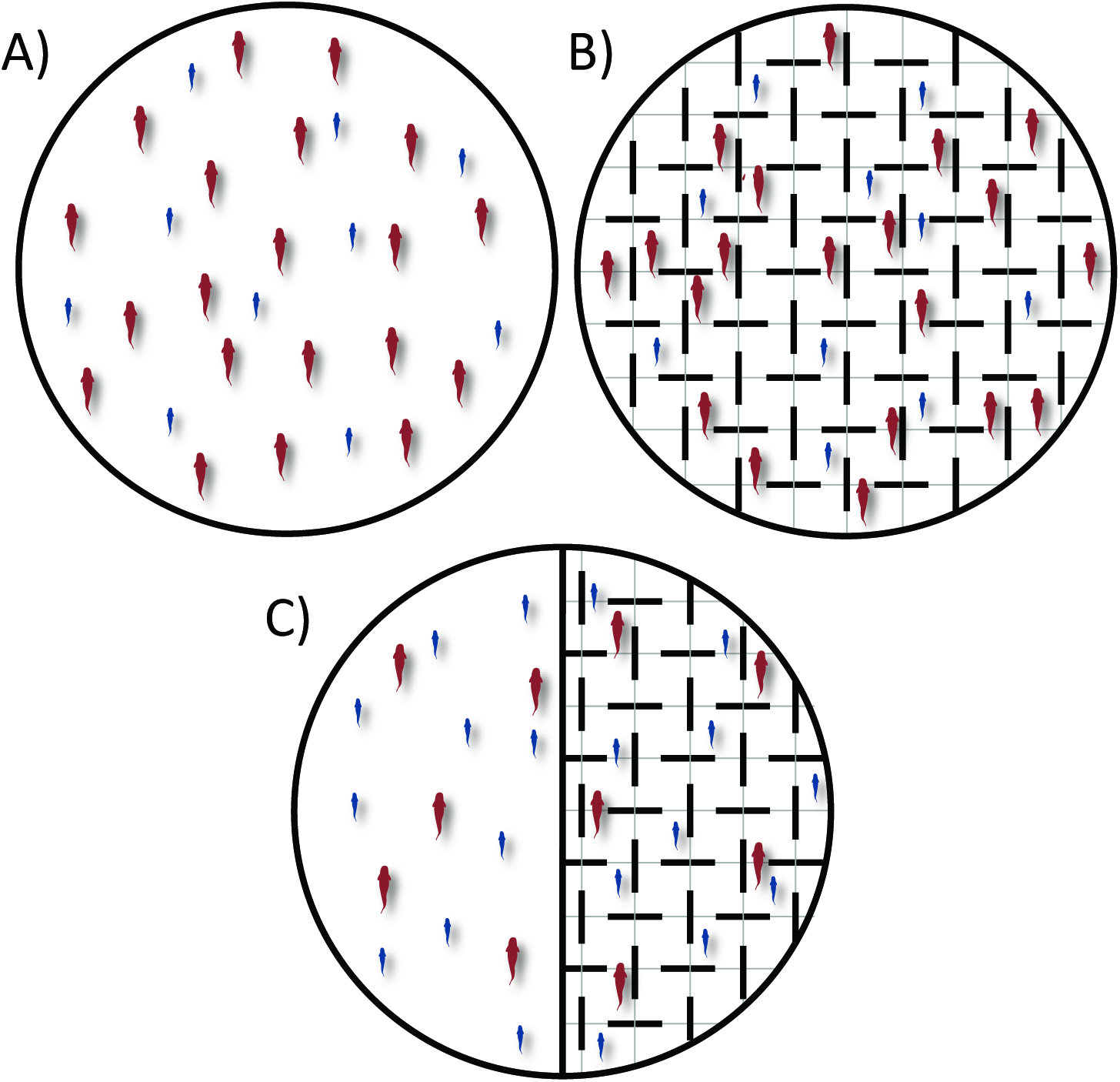
S1 Schematic of experimental ponds A) female biased simple habitat, B) female-biased complex habitat, C) male-biased simple (left) and complex (right). Ponds were 1m in diameter and water depth was 15cm.

DNA samples were sent to the commercial genotyping service Diversity Arrays This company has developed a widely used technique called DArTseq™. DArTseq™ represents a combination of DArT complexity reduction methods and next generation sequencing platforms (Kilian et al, 2012; Courtois et al, 2013; Cruz et al. 2013; Raman et al. 2014;). It is a new implementation of sequencing complexity reduced representations (Altshuler et al, 2000) and more recent applications of this concept on next generation sequencing platforms (Baird et al, 2008; Elshire et al, 2011). The technology is optimized by selecting the most appropriate complexity reduction method based on both the size of the representation and the genome fraction selected for assays. Four methods of complexity reduction were tested in *Gambusia* and the PstI-HpaII method was selected. DNA samples were processed in digestion/ligation reactions principally as per Kilian et al (2012), but replacing a single PstI-compatible adaptor with two different adaptors corresponding to two different Restriction Enzyme (RE) overhangs. The PstI-compatible adapter was designed to include Illumina flowcell attachment sequence, sequencing primer sequence and “staggered”, varying length barcode region, similar to the sequence reported by Elshire et al (2011). The reverse adapter contained flowcell attachment region and HpaII-compatible overhang sequence. Only “mixed fragments” (Pstl-Hpall) were effectively amplified in 30 rounds of PCR using the following reaction conditions: 1. 94° C for 1 min; 2. 30 cycles of 94° C for 20 sec, 58° C for 30 sec, 72° C for 45 sec; 3. 72 C for 7 min. After PCR equimolar amounts of amplification products from each sample of the 96-well microtiter plate were bulked and applied to c-Bot (Illumina) bridge PCR followed by sequencing on Illumina Hiseq2500. The sequencing (single read) was run for 77 cycles.

Sequences generated from each lane were processed using proprietary DArT analytical pipelines. In the primary pipeline the fastq files were first processed to filter out poor quality sequences, applying more stringent selection criteria to the barcode region than the rest of the sequence. In that way the assignments of the sequences to specific samples carried in the “barcode split” step are very reliable. Approximately 2 500 000 (±7%) sequences per barcode/sample are used in marker calling in routine DArTseq assay, but we applied a more cost effective version using 1 300 000 per sample). Finally, identical sequences were collapsed into “fastqcall files” used in the secondary pipeline for DArT PL’s proprietary SNP and SilicoDArT (presence/absence of restriction fragments in representation) calling algorithms (DArTsoft14). This clean-up process resulted in a comprehensive data set of approximately 3171 SNPs with an average call rate of 97.7% and a reproducibility rate of 99.3%.

